# Behavior of *Caenorhabditis elegans* in a nicotine gradient modulated by food

**DOI:** 10.1101/079699

**Authors:** Robert Sobkowiak, Piotr Kaczmarek, Mateusz Kowalski, Rafał Kabaciński, Andrzej Lesicki

**Affiliations:** Department of Cell Biology, Adam Mickiewicz University, Faculty of Biology ul. Umultowska 89, 61-614 Poznań, Poland; Poznan University of Technology, Faculty of Electrical Engineering, Institute of Control and Information Engineering, ul. Piotrowo 3a, 60-965 Poznań, Poland

**Keywords:** Nicotine gradient, Nicotine exposure, Behavior, Body size, Food intake, *Caenorhabditis elegans*

## Abstract

Nicotine decreases food intake, and smokers often report that they smoke to control their weight. To see whether similar phenomena could be observed in the model organism *Caenorhabditis elegans*, we challenged drug-naïve nematodes with a chronic low (0.01 mM) and high (1 mM) nicotine concentration for 55 h (from hatching to adulthood). After that, we recorded changes in their behavior in a nicotine gradient, where they could choose a desired nicotine concentration. By using a combination of behavioral and morphometric methods, we found that both nicotine and food modulate worm behavior. In the presence of food the nematodes adapted to the low nicotine concentration, when placed in the gradient, chose a similar nicotine concentration like *C. elegans* adapted to the high nicotine concentration. However, in the absence of food, the nematodes adapted to the low nicotine concentration, when placed in the gradient of this alkaloid, chose a similar nicotine concentration like naïve worms. The nematodes growing up in the presence of high concentrations of nicotine had a statistically smaller body size, compared to the control condition, and the presence of food did not cause any enhanced slowing movement. These results provide a platform for more detailed molecular and cellular studies of nicotine addiction and food intake in this model organism.

## 1 Introduction

Tobacco smoking is the largest single preventable cause of many chronic diseases and death (WHO, 2012). Most of the tobacco cigarette's toxicity is related to the combustion process. Although tobacco smoking has declined since the 1950s, the use of e-cigarettes has increased, attracting both former tobacco smokers and never smokers. Regardless of nicotine source, method for delivering, and other concomitant substances, nicotine is the most biologically active ingredient of seriously harmful tobacco smoke and the potentially harmful e-cigarette aerosol (Grana et al., 2014; Schuller, 2009; Sobkowiak and Lesicki, 2011).

Some smokers report that they smoke as a method of weight control (Nichter et al., 2004). Indeed, smokers have a notably lower body mass index than nonsmokers and gain weight when they quit (Filozof et al., 2004; Jo et al., 2002; Mineur et al., 2011). These effects on body weight have been attributed to nicotine in tobacco, because nicotine decreases food intake in animal models. In humans, nicotine has some effects on peripheral energy metabolism (Filozof et al., 2004; Jo et al., 2002) and this was also observed in the model nematode *Caenorhabditis elegans* (Sobkowiak et al., 2016).

Nicotine competes with acetylcholine for binding to specific membrane receptors, so-called nicotinic cholinergic receptors (nAChRs). They are widely expressed in the nervous system and skeletal muscle. Nicotinic receptors are also present in many cell types, e.g. epithelial, blood, fat, and cancer cells (Liu et al., 2004; Sobkowiak and Lesicki, 2011). Also nicotine metabolites have significant effects on the body (Benowitz et al., 2009; Sobkowiak and Lesicki, 2013). Early nicotine exposure has been associated with many long-term consequences that include neuroanatomical alterations as well as behavioral and cognitive deficits (Rose et al., 2013). Moreover, nicotine is a highly addictive substance with negative effects on animal and human brain development, which is still ongoing in adolescence (Benowitz, 2010; Grana et al., 2014).

Locomotion reflects the integration of many aspects of both the environment and the internal state of the worm, and therefore can be a sensitive measure of behavioral state after nicotine application. As an animal travels through its environment, its nervous system detects sensory cues, evaluates them based on context and the experience of the animal, and converts the information into adaptive movement. *C. elegans* has behavioral states that are evident as long-term locomotor patterns that differ in well-fed or starved animals. When feeding on a bacterial lawn, *C. elegans* switches between 2 behavioral patterns. It spends 80% of its time dwelling, i.e. feeding on bacteria while moving slowly and staying in a restricted area. At rare intervals, however, it switches into an alternative behavioral state called roaming, which involves rapid locomotion across the lawn. When removed from the bacterial lawn, *C. elegans* switches to a behavioral state in which it moves rapidly and reverses frequently. This strategy is called pivoting, area-restricted search, or local search, and has the same basic run-and-pirouette components as directed chemotaxis. Over the next 30 min, reversals are suppressed, shifting the animals to another behavioral state, called traveling or dispersal (Bargmann, 2006 and the papers cited therein).

Worms navigate to favorable conditions by chemotaxis and aerotaxis (Gray et al., 2005). *C. elegans* shows chemotaxis to various odorants and water-soluble chemoattractants, like nicotine, by sensing them mainly with sensory neurons whose sensory endings are located in amphid sensory organs at the anterior tip of the animal. The strategy of chemotaxis in this organism was previously studied by Pierce-Shimomura et al. (1999), who found the “pirouette strategy” for chemotaxis to water-soluble chemoattractants. Iino and Yoshida (2009) reported the discovery of a second mechanism for chemotaxis, called the weathervane mechanism. In that strategy, the animals respond to a spatial gradient of the chemoattractant and gradually curve toward a higher concentration of the chemical.

*Caenorhabditis elegans* is a powerful genetic model for studying various questions in neurobiology, including nicotine dependence. For example, it has been reported that *C. elegans* responds to nicotine in both a concentration-dependent and a time-dependent manner, and shows a wide range of behavioral responses to nicotine (Feng et al., 2006; Sobkowiak et al., 2011). These include a continuous higher locomotion speed (acute response), tolerance, withdrawal, and sensitization via nAChR (Feng et al., 2006). Dual effects of nicotine on locomotion speed, which are dependent on differences in its dosage and treatment duration, have also been revealed (Sobkowiak et al., 2011). Moreover, *C. elegans* is capable of navigating up a nicotine gradient (Sellings et al., 2013).

In the current study we observed spontaneous behavior in the presence or absence of food as well as a nicotine gradient to describe the effects of low and high nicotine concentration exposure during larval development in adult *C. elegans*.

## 2 Materials and methods

### 2.1 *Caenorhabditis elegans* maintenance

All tests were performed on the wild-type Bristol N2 strain of *C. elegans* obtained from the Caenorhabditis Genetics Center (CGC) at the University of Minnesota (Duluth, Minnesota, USA). Standard methods were used for the maintenance and manipulation of strains (Stiernagle, 2006). Nematodes were kept at 22°C on nematode growth medium (NGM) agar plates seeded with *Escherichia coli* strain OP50 as a source of food according to a standard protocol (Brenner, 1974). Chunks of mixed-stage starved worms were transferred onto enriched nematode growth (ENG) plates (50 mm in diameter) seeded with bacterial food (*E coli* OP50), and the worms were allowed to grow for about 5 days at 22°C (Figure 1a).

**Figure 1.**
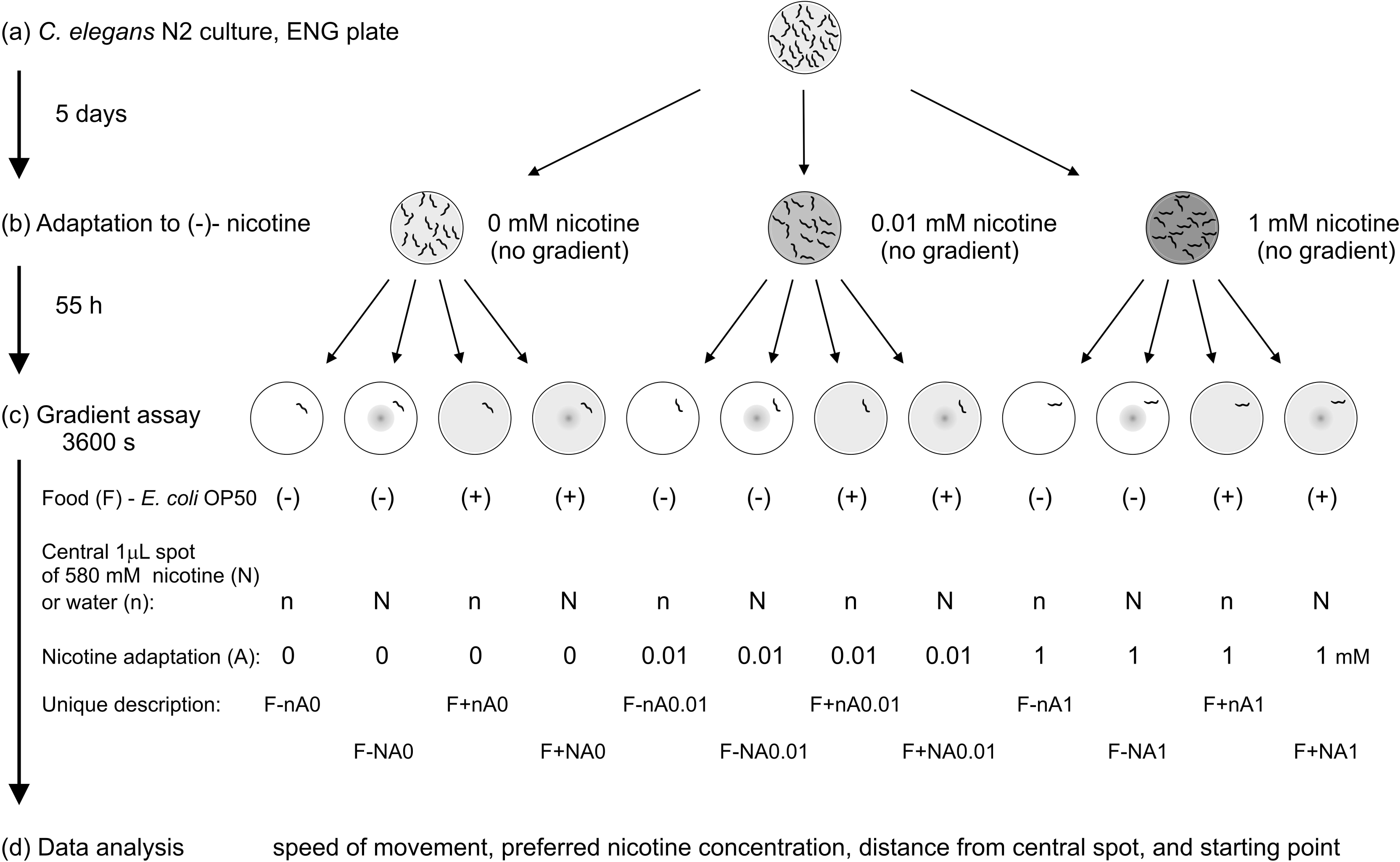
Outline of experimental procedures. (a) *Incubation*. *Caenorhabditis elegans* were grown for 5 days at 22°C on ENG medium plates spread with *E. coli.* (b) *Long-term nicotine adaptation*. Adult worms were transferred to NGM plates seeded with food, by transferring of chunks of the medium with nematodes onto control plates (0 mM nicotine) and adaptation plates containing 0.01 mM nicotine (low nicotine concentration) or 1 mM nicotine (high nicotine concentration). All adult worms after 3 h were removed and only the laid eggs were left. Incubation at 22°C for 55 h. (c) *Transfer of single adult nematodes onto gradient assay plates*. Duration of the experiment: 3600 s. White plate = no food; uniform gray plate = presence of food; peak at the plate center = application point of nicotine at the beginning of the experiment; no peak at the plate center = control plates (with water instead of nicotine). (d) *Data analysis of used behavioral endpoints measured in the tests*.

### 2.2 Chronic exposure of *C. elegans* to uniform nicotine concentration

(-)-Nicotine (free base) was obtained from Sigma-Aldrich. The drug was added directly to the NGM medium and allowed to diffuse throughout the medium for 48–72 h to ensure a uniform concentration.

To minimize interaction between *E. coli* and nicotine, 10 drops of liquid culture of *E. coli* OP50 were added just before chronic exposure of *C. elegans* to nicotine on agar plates. Bacteria were evenly distributed by a spreader on the surface of the NGM medium, and the plates with the lids removed were left to dry for about 10 min. The short drying of the agar surface enabled the nematodes free movement and proper nutrition.

When the culture of nematodes contained mostly adults able to lay eggs, chunks of the agar with nematodes were transferred onto the plates for chronic exposure to a uniform nicotine concentration (0.01 M or 1 mM, Figure 1b). In the controls, the chunks were transferred to nicotine-free NGM plates. To synchronize the worms, parent worms were allowed to lay eggs for 3 h and next the adults were removed. The remaining eggs were incubated for 55 h at 22°C to adulthood (Altun and Hall, 2009). The dosages were selected based on our preliminary study as well previous reports (Feng et al., 2006; Matta et al., 2007; Sobkowiak et al., 2011; Waggoner et al., 2000), in which nicotine treatment had a biphasic response.

### 2.3 Measurement of worm size

The measurements of worm size and the behavioral nicotine gradient experiments were performed using adult hermaphrodites, which were kept in the presence of nicotine from hatching to adulthood. Body size of adult worms was estimated in control conditions (0 mM nicotine) and after 55 h exposure to 0.01 and 1 mM nicotine. To measure body length, volume, and surface area, the worms were put on an NGM plate, and pictures were taken using a stereomicroscope and analyzed using WormSizer, as previously described (Moore et al., 2013). WormSizer is open-source software that is useful for detecting relatively subtle phenotypes and morphological changes that may have been difficult to assess upon visual inspection. Eight biological replicates, typically including 12 individuals per experimental variant, were analyzed.

### 2.4 Behavioral nicotine gradient assay

In all the experiments we used young adult worms exposed to 0 mM (control), 0.01 mM, and 1 mM nicotine. After 55 h of nicotine exposure, single young adult hermaphrodites from each adaptation variant were transferred to assay plates containing NGM medium with 4 treatment variants (Figure 1c):

1. no food, no radial Gaussian-shaped nicotine gradient (F-n);
2. no food, a radial Gaussian-shaped nicotine gradient (F-N);
3. spatially uniform concentration of food, no radial Gaussian-shaped nicotine gradient (F+n);
4. spatially uniform concentration of food, a radial Gaussian-shaped nicotine gradient (F+N).

The behavioral nicotine gradient assays were performed in standard Petri dishes (inner diameter 92 mm) containing 5.8 mL of NGM. In variants 3 and 4, NGM agar was seeded with 3 drops of the same medium with *E. coli* OP50, while in variants 1 and 2 the plates were seeded with 3 drops of sterile medium without *E. coli* (i.e. without food), and allowed to desiccate for about 10 min. The radial Gaussian-shaped nicotine gradient was formed by adding 1 μL of 580 mM nicotine solution in water at the center of the plate just before the tracking. Nicotine concentration was estimated in the range of 0-580 mM, according to the diffusion equation for a point bolus in a thin slab (Crank, 1975) (Figure 2b).

**Figure 2.**
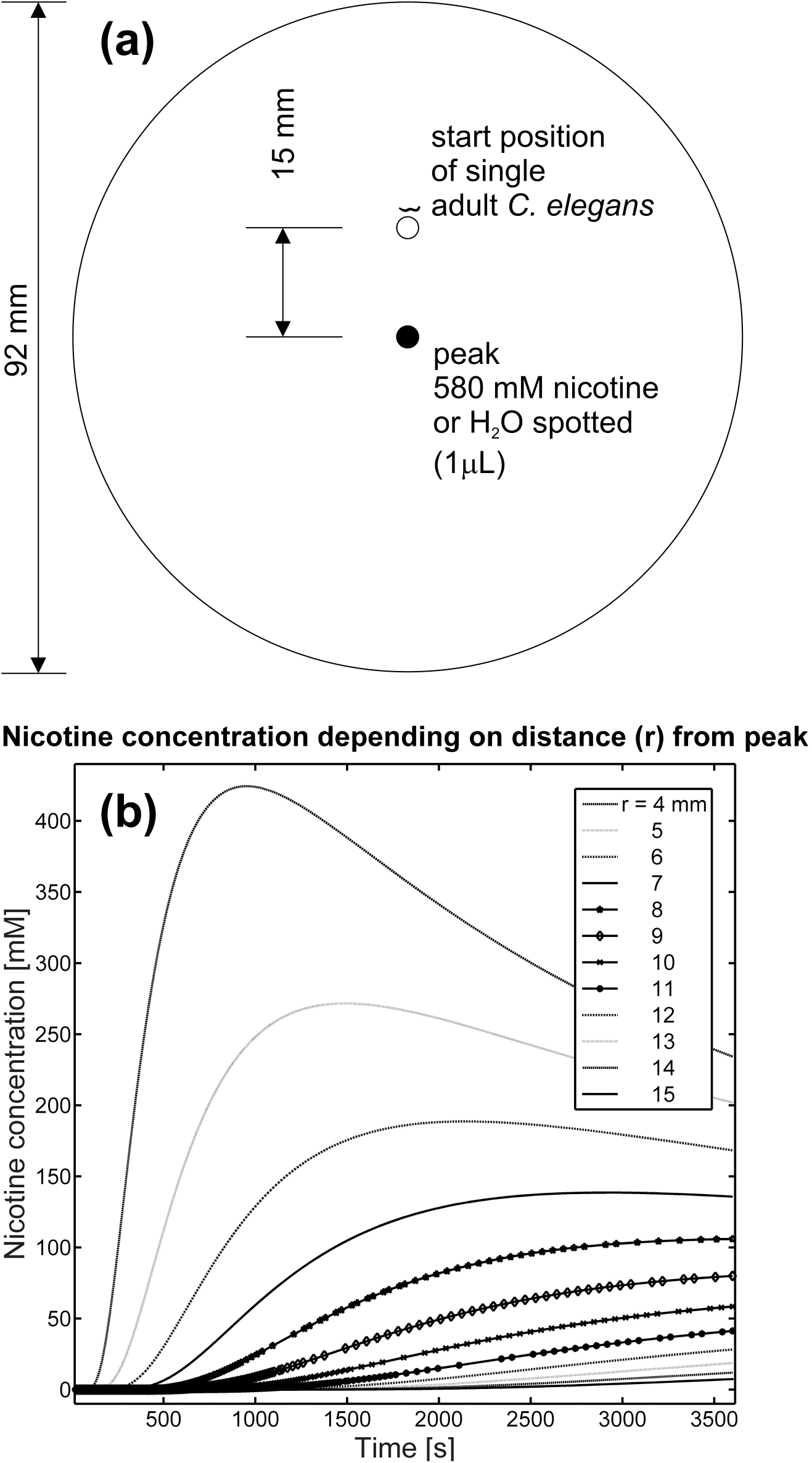
**(a)**Configuration of the plate for nicotine gradient assay of *Caenorhabditis elegans*. One worm was placed in a 1-μL drop of water and thus trapped for several dozen seconds in the start position of a Petri dish containing 5.8 mL of NGM medium. Immediately after the worm was released, nicotine was spotted into the center of the plate (peak). Water was applied instead of nicotine in control variants. **(b)** Theoretical distribution of nicotine concentration depending on the distance (*r* = radius) from the peak. The gradient was formed by placing 1μL of 580 mM (-)-nicotine at the center of the plate at the beginning of experiment. Nicotine appears to move smoothly and systematically from high-concentration areas to low-concentration areas, following Fick's laws. Concentration estimates were made as described in section 2.4 (see the equation)

For each assay plate, time was recorded for estimation of nicotine concentration during the assay. At each time point of the assay, the concentration of nicotine (mM) at the position of the worm was estimated according to the solution of the diffusion equation (Crank, 1975) for a point bolus in a cylindrical, aqueous volume having the same dimensions as the agar in the assay plate (diameter 9.2 cm; depth 0.1 cm). Exact nicotine concentration (in mM) was calculated by the solution of Fick's equation for 2-dimensional diffusion with no border (Crank, 1975; Iino and Yoshida, 2009) (Figure 3).

**Figure 3.**
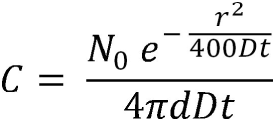
The Fick's equation for 2-dimensional diffusion with no border: *C* = nicotine concentration [mM]; *N*_0_ = 0.58 [mmole] is the number of moles of nicotine spotted (1 μL of 580 mM nicotine contains 0.58 mmoles of nicotine); *D* = 0.000 000 42 [cm^2^/s] is the diffusion coefficient of nicotine (estimated basing on Sellings et al., 2013); *d* = 0.1 [cm] is the depth of the agar; *t* = is the time [s] after spotting nicotine at the center of the plate; *r* = is the distance [cm] between the peak of the gradient and the location of the animal (Crank, 1975; Iino and Yoshida, 2009).

To study the behavior of worms, we used an automated tracking system to follow individual young adults crawling on NGM plates (Kowalski et al., 2014). Single young adults (aged ~ 55 h) were manually picked off an adaptation plate (Figure 1b) and placed using a platinum pick in a 1-μL droplet of water on the agar surface of the gradient assay plate (Figure 1c), 15 mm from the plate center (Figure 2a). Putting a worm in a droplet of water is an effective method for rapid transferring of a single animal without scratching the agar surface (important for obtaining high-contrast videos and perfect tracking) and for testing the condition of the worm (injured nematodes, which could not properly swim in a drop of water, were rejected). The surface tension of the drop of water prevented the nematode from creeping out. Next, the plate was placed in a device for tracking nematodes. Observation of the nematode allowed us to notice the moment when the 1-μL drop disappeared by evaporation/absorption in agar and released the worm, which could then freely move on the plate. At that moment, in the center of the plate, 1 μL of 580 mM nicotine or 1 μL of H_2_O was spotted (Figures 1c and 2a). Worm tracking began no more than 10 s after the application of nicotine or water in the center of the plate. Each worm was tracked for 3600 s or until it reached the edge of the plate.

### 2.5 Worm tracking system

Custom worm tracker software (WormSpy) was used to move the camera automatically to re-center the worm under the field of view during recording (Kowalski et al., 2014). The automated tracking system comprises a stereomicroscope (Olympus SZ11), a modified (with unscrewed lens) web camera (Logitech QuickCam Pro 9000) with 640×480 resolution to acquire worm videos, and a desktop PC running under Windows 7. The tracking system located the worm's centroid (defined as the geometrical center of the smallest rectangle that could be drawn around the worm) and recorded its *x* and *y* coordinates with a sampling rate of 2 s^−1^. When a worm neared the edge of the field of view, the tracking system automatically re-centered the worm by moving the stage and recorded the distance that the stage was moved. We reduced the variation in sampling rate as a consequence of the small differences in the time it took to re-center the worm and the need to take data only when the stage was stationary by developing a simultaneous localization and tracking method for a worm tracking system (Kowalski et al., 2014). The spatiotemporal track of each worm was reconstructed from the record of centroid locations and camera displacements. The instantaneous speed and trajectory were computed using the displacement of the centroid in successive samples. The tracking system recorded the worm's position, speed, distance from the center of the plate and from the starting point, estimated nicotine concentration in the surroundings the worm, and trajectory at 0.5-s intervals. Individual worms moved away from their starting location, leaving complex tracks. Video recordings were carried out at room temperature (22°C). Five independent experiments were performed per dosage group.

All the experimental procedures presented in this paper were in compliance with the European Communities Council Directive of 24 November 1986 (86/609/EEC).

### 2.6 Data analysis and statistical analysis

The measurements from experiments were pooled for each treatment variant and the median values were calculated. The data were not normally distributed, as determined by the Shapiro-Wilk *W*-test and Kolmogorov-Smirnov & Lilliefors method. Due to this, the Kruskal–Wallis tests followed by Dunn's multiple comparison post-hoc tests were performed. Statistical significance was considered at *p* < 0.05. The calculations and graphs were done by using Statistica software (StatSoft, Inc., Tulsa, Oklahoma, USA).

Worm body size data are presented as mean±S.D. of at least 8 independent experiments. One-way ANOVA followed by Scheffe's test were performed to determine statistical differences between groups with the aid of Statistica (StatSoft software). Significance was set at *p* < 0.05.

## 3 Results

### 3.1 Body size of worms

Nicotine affected the body size of nematodes. Worms challenged from hatching to adulthood (55 h, 22^o^C) with 1 mM nicotine were smaller than in control conditions (Figure 4). The worms adapted to the high nicotine concentration (1 mM) were about 5% shorter (Figure 4a), with about 12% smaller surface (Figure 4b), and about 22% smaller body volume, as compared to control nematodes (Figure 4c). The nematodes adapted to a low nicotine concentration (0.01 mM) were on average about 4.5% longer (Figure 4a), with about 5.5% smaller volume (Figure 4c), as compared to control nematodes. However, there was no statistically significant difference between these 2 groups.

**Figure 4.**
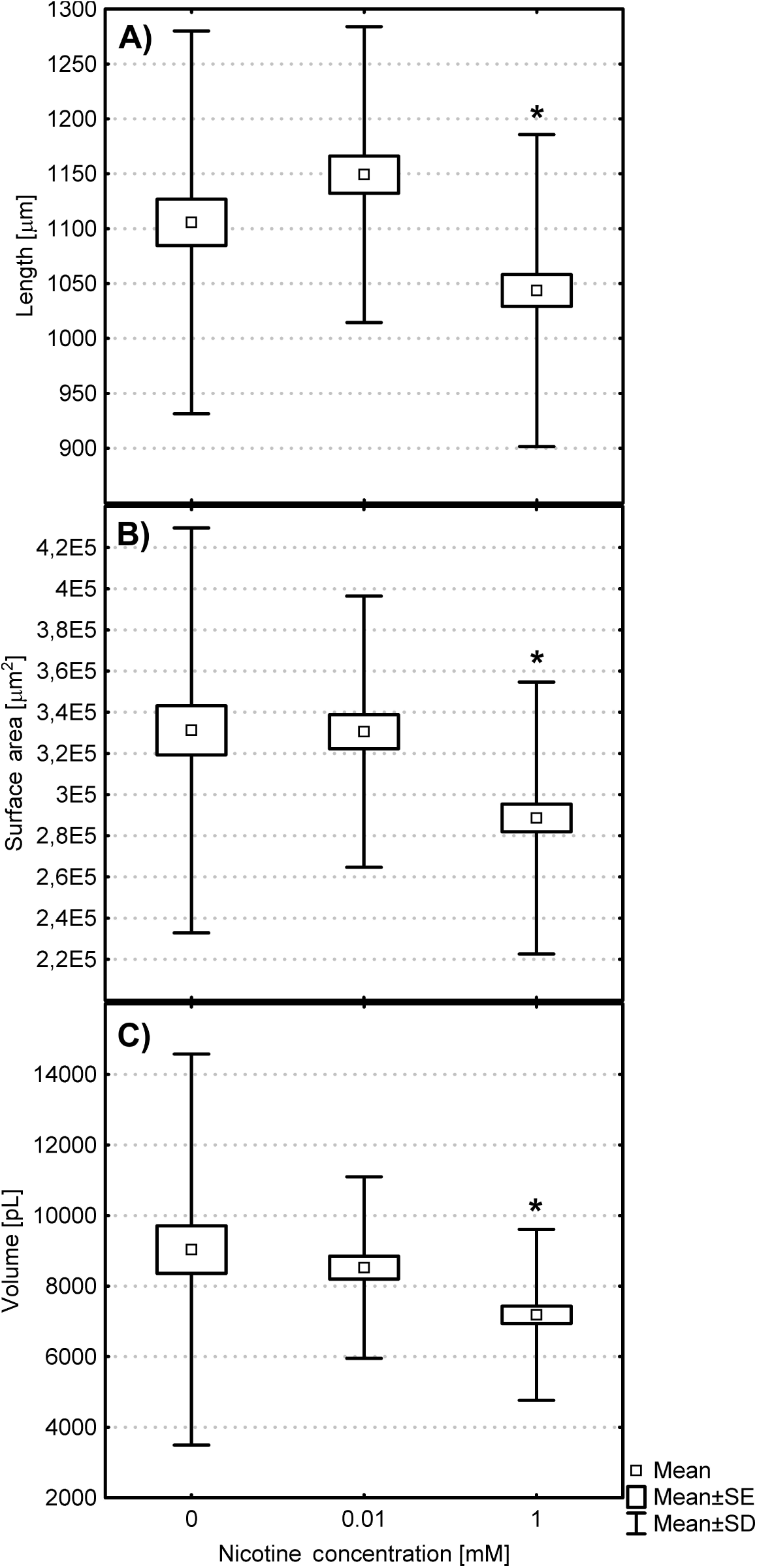
Differences in body size of *Caenorhabditis elegans* after 55 h of growth in control conditions (0 mM nicotine) and in the presence of 0.01 mM nicotine and 1 mM nicotine (*\* p* < 0.05, Scheffe test, *N* = 96 at least).

### 3.2 Average values of factors describing worm behavior

#### 3.2.1 Average speed of movement

The average speed of movement of well-fed *C. elegans* worms (Figure 5a) was the fastest in the absence of food (F-nA0), and their locomotion slowed down when food was present (F+). A similar effect of slowing the movement, but also in the absence of food, was evoked by the gradient of nicotine (F-N).

**Figure 5.**
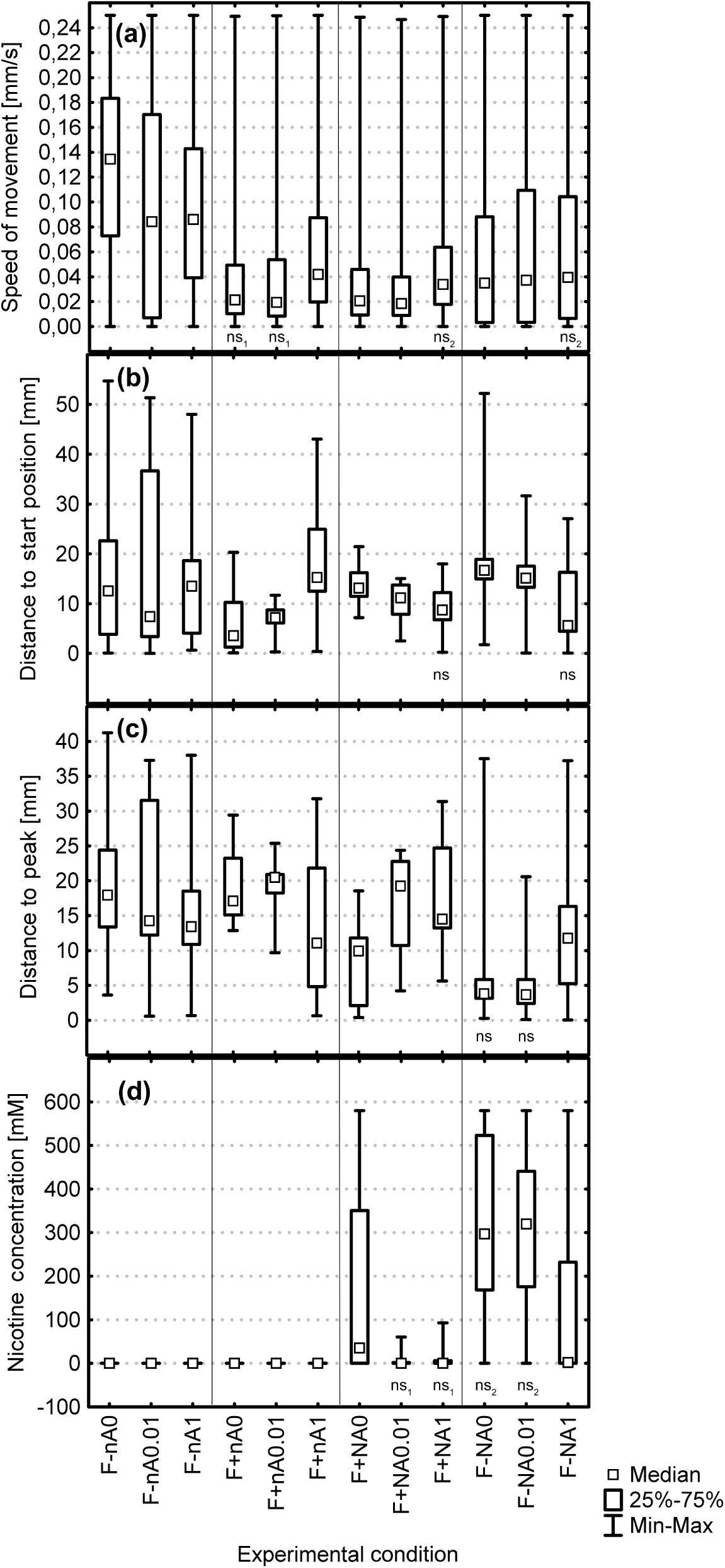
Average values of factors describing the behavior of *Caenorhabditis elegans*: (a) speed of movement; (b) distance from the start position; (c) distance from the peak; (d) preferred nicotine concentration; “ns” denoted no statistically significant differences between experimental conditions. Other groups were statistically significantly different from each other (*p* < 0.05, Kruskal-Wallis ANOVA by ranks test, data pooled from 5 independent experiments, *N* = 14 370).

In the presence of food on the nicotine-free Petri dish, there was no statistically significant difference in locomotion speed between the nematodes having the first contact with this alkaloid (naïve worms) and nematodes adapted to 0.01 mM nicotine (Figure 5a, F+nA0 vs. F+nA0.01). In the presence of food and without nicotine in the center of the assay plate, the worms adapted to 0 mM nicotine (F+nA0) and those adapted to 0.01 mM nicotine (F+nA0.01) moved with the same speed (no significant differences, Figure 5a).

The worms adapted to 1 mM nicotine in the presence of nicotine gradient moved with the same speed independently of the presence or absence of food (Figure 5a, no significant differences between F+NA1 and F-NA1). It is noteworthy that the nematodes adapted to 1 mM nicotine (F+nA1) in the presence of food unexpectedly moved farthest from the start position and the presence of food appeared to stimulate the process.

#### 3.2.2 Average distance from the start position

Nematodes moved away from the starting point in a range of 3 mm (F+KA0) to 18 mm (F-NA0) (Figure 5b). The presence of food in the absence of nicotine gradient evoked in naïve worms a tendency to keep close to the start position (Figure 5b, F-nA0 vs. F+nA0). Median value of the distance from the starting point for the worms adapted to 0.01 mM nicotine was almost the same but the range of 25%-75% narrowed (Figure 5b, F-nA0.01 vs. F+nA0.01).

In the presence of food we observed a linear tendency with a significantly (*p* < 0.05, Kruskal-Wallis ANOVA by ranks test) increased distance of worms from the starting point (Figure 5b, F+nA0 lowest value, F+nA0.01, F+nA1 highest values).

We observed an opposite tendency when a gradient of nicotine appeared on the plate. Independently of the presence or absence of food, nicotine extorted a decrease in the distance from the start position in line with the increase in nicotine concentration to which the worms were earlier adapted (Figure 5b, F+NA0, F+NA0.01, F+NA1). Interestingly, we revealed no statistical difference in average distance from the start position between the worms adapted to 1 mM nicotine in the presence of food and absence of food (Figure 5b, F+NA1 vs. F-NA1)

#### 3.2.3 Average distance from the peak

The peak was the central point where nicotine or water was applied on the gradient assay plate. The radius in which nematodes were present, measured from the peak to worm location, was in the range from about 4 mm (F-NA0, F-NA0.01) to about 21 mm (F+nA0.01) (median value, Figure 5c). When there was no nicotine in the center of the gradient assay plate, the shortest distance from the peak was observed for the worms adapted to 1 mM nicotine (F+nA1). When the gradient of nicotine appeared and in the absence of food, the worms adapted to 0 mM and 0.01 mM reached the peak and behaved in the same way (Figure 5c F-NA0, F-NA0.01, no statistical difference). The worms adapted to 0.01 mM nicotine in the presence of food were not attracted to the peak and they chose food and kept distance like on the plates without nicotine (Figure 5c, compare median values for F+nA0.01 vs. F+NA0.01 vs. F-NA0.01).

#### 3.2.4 Average nicotine concentration preferred by worms

In our experiment, nematodes could select the concentration of nicotine in which they wanted to stay. Their movements depended on the presence of food and on adaptation to nicotine. In the absence of food, worms preferred on gradient assay plates higher concentrations of nicotine in comparison to the worms placed on plates without food (Figure 5d). The nematodes adapted to the low concentration of nicotine (0.01 mM) behaved differently in comparison to the nematodes adapted to the high concentration of nicotine (1 mM). The worms adapted to 1 mM chose the lowest nicotine concentration independently of the presence or absence of food (Figure 5d, F+NA1 and F-NA1). In the presence of food the worms adapted to 0.01 mM nicotine behaved like the worms adapted to 1 mM nicotine (Figure 5d, F+A0.01 vs. F+NA1, no statistical difference). However, in the absence of food, the worms adapted to 0.01 mM nicotine behaved like the worms adapted to 0 mM nicotine (Figure 5d, F-NA0 vs. F-A0.01, no statistical difference) and chose a very high concentration of nicotine.

### 3.3 Time-dependent response of *C. elegans* to the nicotine gradient

#### 3.3.1 Time-dependent speed of movement

The speed of the worms varied with time (Figure 6). In most cases, we observed a decreasing trend in the speed of their movements, and its frequent oscillations. In control conditions, i.e. in the absence of a gradient of nicotine and food (F-nA0), we observed oscillations of the speed, while the nematodes in variant F-nA0.01 show a downward trend in the speed of movement (Figure 6e). The long period of reduced locomotion activity was observed particularly in control nematodes (Figure 6b, time 500-1100 s) and those adapted to 0.01 mM nicotine (Figure 6e, time 2500-3600 s; Figure 6g, time 700-1400 s).

**Figure 6.**
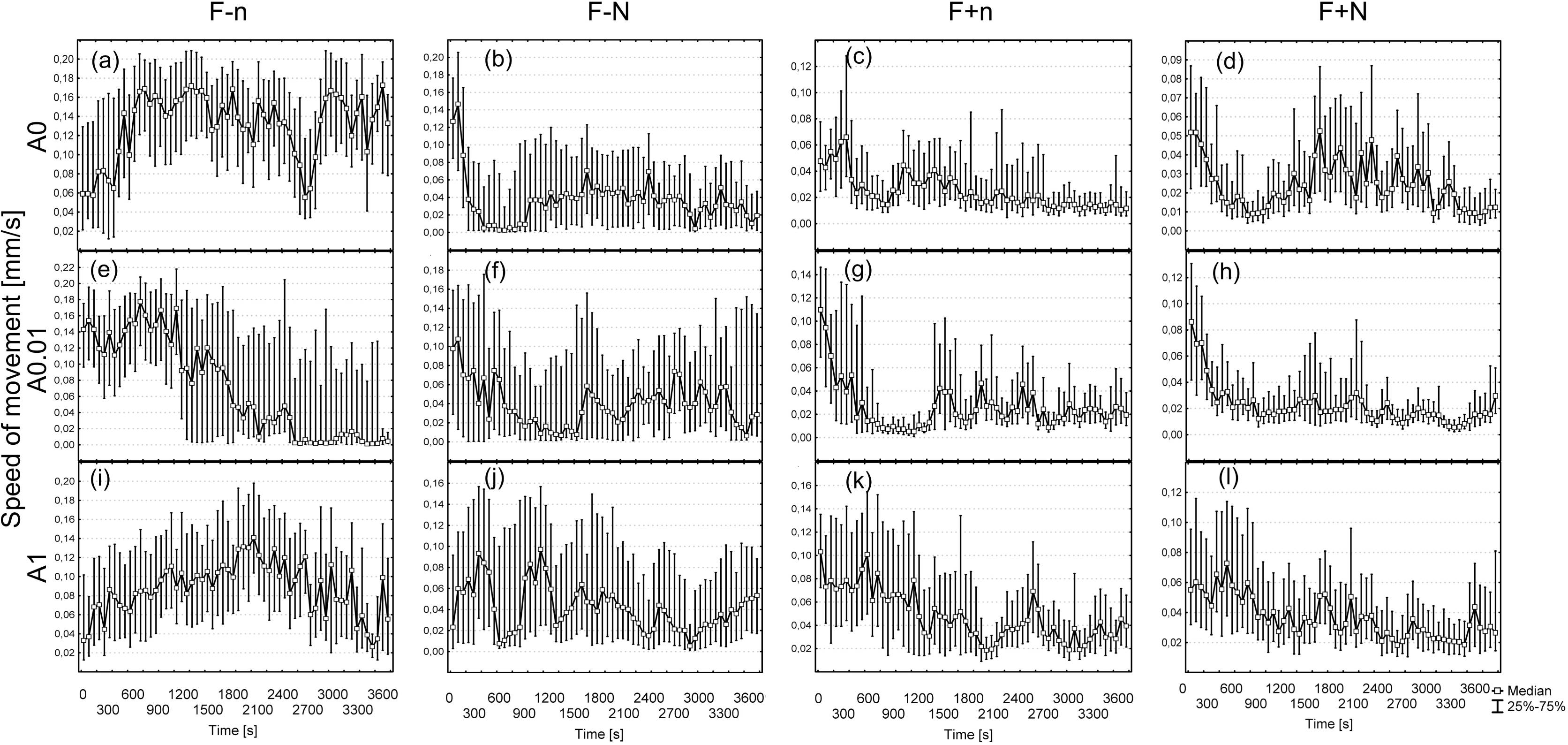
Locomotor activity (centroid speed) of *Caenorhabditis elegans* in the tested experimental variants. The data are medians and 25th and 75th percentiles of 5 pooled experiments. F-/F+ = absence/presence of food, respectively; n = no nicotine gradient; N = presence of nicotine gradient; A0 = naïve worms, which never had contact with nicotine before; A0.01 = worms adapted to 0.01 mM nicotine; A1 = worms adapted to 1 mM nicotine.

#### 3.3.2 Time-dependent changes in distance from the start position

Figure 7 shows the variation in the distance from the start position of worms (measured from the start position to the actual worm location). Generally during the experiment the nematodes moved away from the starting point. In some experimental conditions, nematodes remained at a relatively stable distance from the start position (Figure 7b, f, k). This applied especially to the worms that never had contact with nicotine before, and were transferred to a Petri dish with bacteria and no nicotine gradient (Figure 7c, F+nA0). Those worms tended to move away from the starting point about 1200 s after application (Figure 7c). Farthest mean distance from the start position was recorded for the nematodes adapted to 0.01 mM nicotine, on the plate without nicotine and bacteria (variant F-nA0.01, Figure 7e).

**Figure 7.**
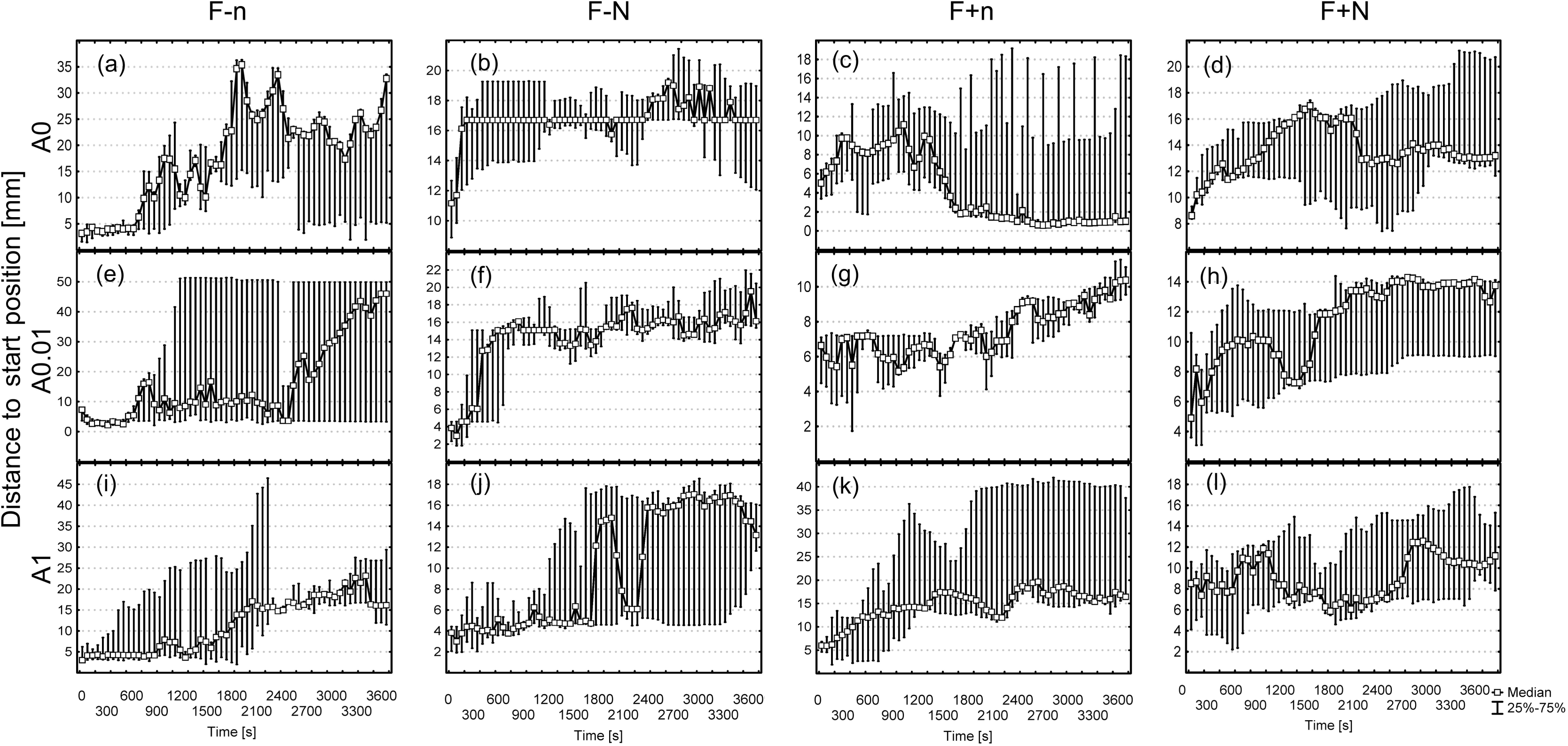
Distance from the start position of *Caenorhabditis elegans* in the tested experimental variants. The data are medians and 25th and 75th percentiles of 5 pooled experiments. F-/F+ = absence/presence of food, respectively; n = no nicotine gradient; N = presence of nicotine gradient; A0 = naïve worms, which never had contact with nicotine before; A0.01 = worms adapted to 0.01 mM nicotine; A1 = worms adapted to 1 mM nicotine.

#### 3.3.3 Time-dependent changes in distance from the peak

The distance between worms and the peak varied with time (Figure 8), but was exceptionally constant for the nematodes adapted to 1 mM nicotine and placed on Petri dishes with food and nicotine gradient (F+NA1, Figure 8i). Interesting behavior of nematodes was observed in variants F-NA0 and F-NA0.01 (Figure 8b, f), where almost immediately after application they moved towards the central peak of nicotine.

**Figure 8.**
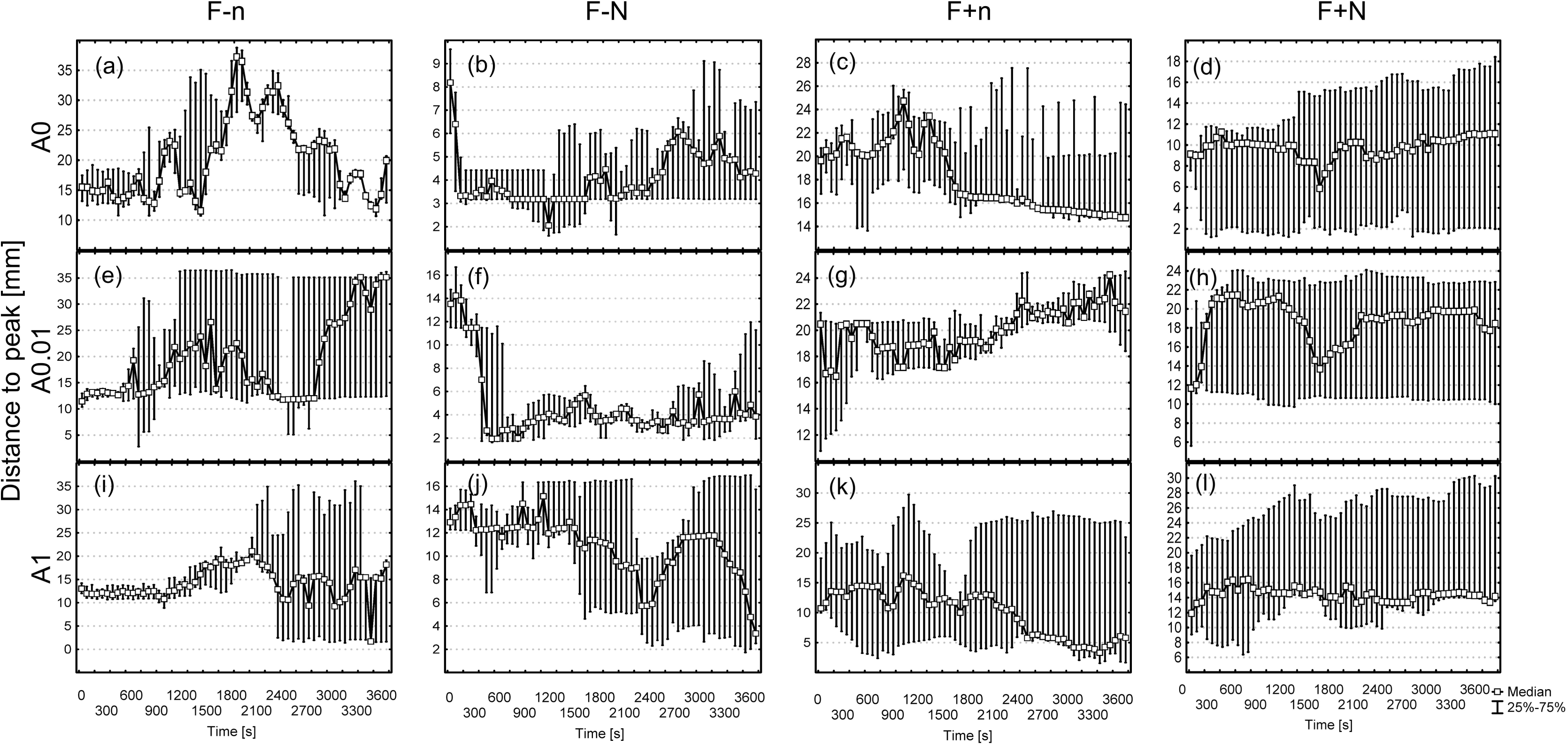
Time-dependent changes in distance from the peak of *Caenorhabditis elegans* in the tested experimental variants. The data are medians and 25th and 75th percentiles of 5 pooled experiments. F-/F+ = absence/presence of food, respectively; n = no nicotine gradient; N = presence of nicotine gradient; A0 = naïve worms, which never had contact with nicotine before; A0.01 = worms adapted to 0.01 mM nicotine; A1 = worms adapted to 1 mM nicotine.

#### 3.3.4 Time-dependent changes in preferred concentration of nicotine

In a radial gradient of nicotine, control wild-type nematodes and those adapted to 0.01 mM nicotine usually reached the gradient peak by moving almost directly up the gradient (Figure 9a, c, F-NA0 and F-NA0.01). The preference for higher concentrations appeared about 300 s after the application of nicotine in the center of the Petri plate. However, the nematodes adapted to 0.01 mM nicotine, when placed on plates with food, avoided nicotine for the first 900 s of experiment (Figure 9d, time 0-1000s). Nematodes adapted to 1mM nicotine avoided nicotine for the first 1500 s of the experiment (Figure 9e, f, time 0-1000s).

**Figure 9.**
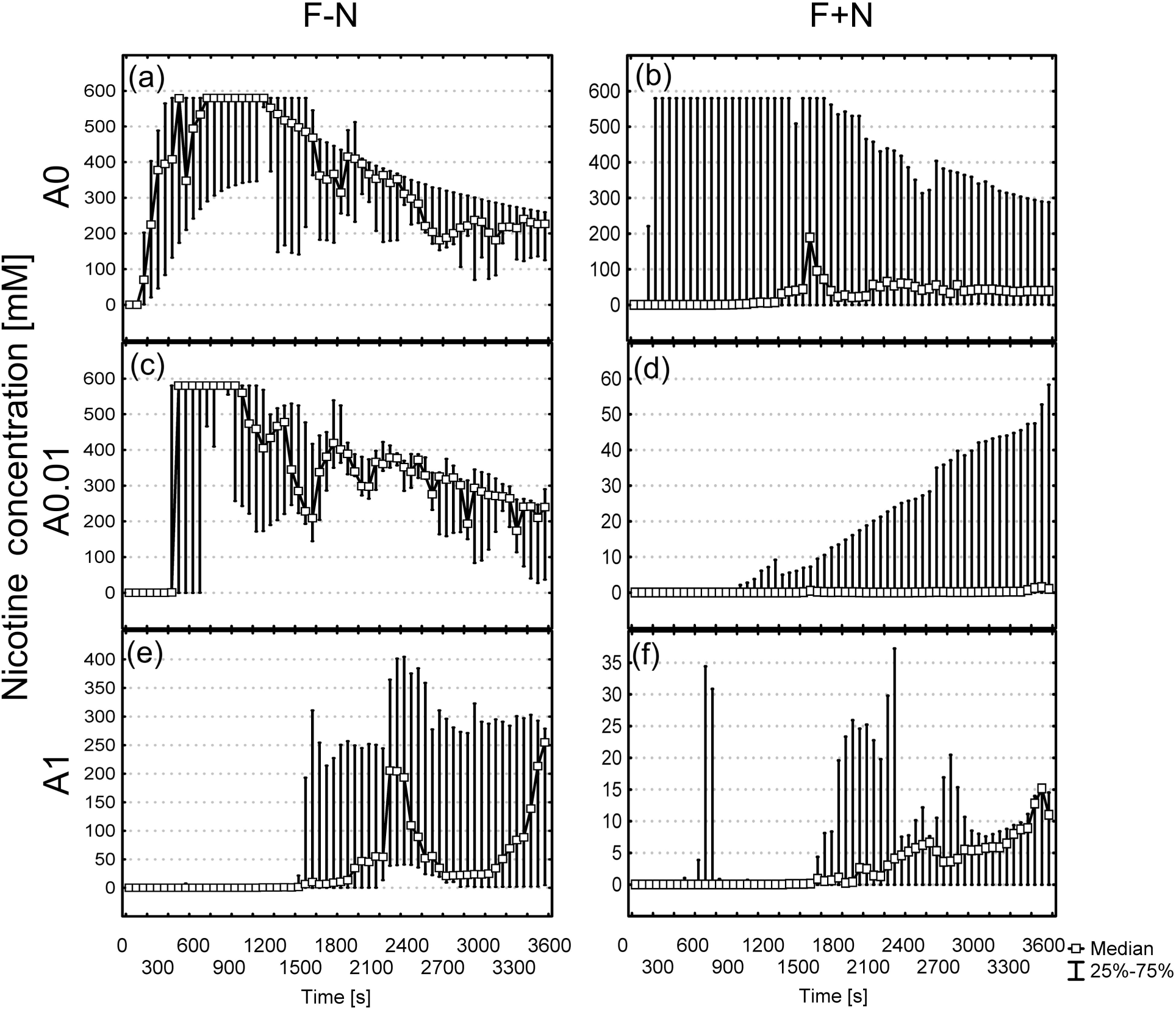
Time-dependent changes in nicotine concentration preferred by *Caenorhabditis elegans* in the tested experimental variants. The data are medians and 25th and 75th percentiles of 5 experiments. F-/F+ = absence/presence of food, respectively; n = no nicotine gradient; N = presence of nicotine gradient; A0 = naïve worms, which never had contact with nicotine before; A0.01 = worms adapted to 0.01 mM nicotine; A1 = worms adapted to 1 mM nicotine.

## 4 Discussion

In the experiments we used 3 groups of nematodes: naïve worms (which never had contact with nicotine before), worms adapted to a low uniform nicotine concentration (0.01 mM) and to a high uniform nicotine concentration (1 mM) (Figure 1b). The adaptation to nicotine lasted from hatching to adulthood, and next we investigated the behavior of these nematodes in a nicotine gradient (Figure 1c). Nicotine was administered in the center of the Petri plate in one bolus dose. Theoretically, the bolus of 1 μL of 580 mM nicotine diluted in 5.8 mL of medium in Petri dish diffuses to final 0.1 mM nicotine concentration and disappearing of nicotine gradient after about 12 h. However, we started our experiments immediately after administration of nicotine onto the Petri dish. In our previous study, in uniform nicotine concentration, within the first 3600 s we observed the largest difference in the behavior of control naïve nematodes and those in the presence of 0.1 mM nicotine (Sobkowiak et al., 2011). In this study, we used a 10-fold lower (0.01 mM) and 10-fold higher nicotine concentration (1 mM) during adaptation of worms. These 2 concentrations were considered “effective dosages”, as our previous report revealed that dosages in this range cause changes in locomotion (Sobkowiak et al., 2011, see Figure 2 in that article).

We performed chemotaxis assays by tracking individual worms in the radial Gaussian-shaped gradients of nicotine. *C. elegans* has a highly developed chemosensory system that enables it to detect a wide variety of volatile (olfactory) and water-soluble (gustatory) cues associated with food and chemicals (Bargmann, 2006). During the experiments, the nematodes were allowed to move freely around the Petri dishes, and able to select the desired nicotine concentration. Sellings et al. (2013) suggest that nicotine acts as a rewarding substance in *C. elegans*. The nematodes approach a point source of nicotine in a time-dependent and concentration-dependent manner. Those authors revealed that wild-type worms climb the nicotine gradient, and suggested that worms exposed to 50 mM nicotine can approach appetitive stimuli, once removed from the nicotine source. This is in line with our result, with the exception of the high nicotine concentration experimental variant. The naïve animals and worms adapted to low nicotine concentration reached the peak of the gradient, defined as a circular region with a radius of 5 mm located at the center of the plate (Figure 2a, Figure 8b, f), and were trapped in the nicotine area when the nicotine concentration exceeded 500 mM at the beginning of experiment (Figure 9a, b, c). The cuticle of *C. elegans* is a significant barrier for drug permeability, thus the internal concentrations in the worm's body fluid is likely to be substantially lower than the dosing medium concentration (Wolf and Heberlein, 2003). However, high concentrations of nicotine induce muscle hypercontraction paralysis (Matsuura et al., 2013; Sobkowiak et al., 2011) and cause rapid transient paralysis of body wall muscles, manifested in an extremely low speed of locomotion (Figure 6a, time 300-900s; Figure 6f, 1200-1600s). This suggests that the nematodes were affected by nicotine toxicity and paralysis.

The naïve animals and those adapted to the low nicotine concentration tested in the nicotine gradient were significantly more unlikely to reach the center when on the Petri dish the food was in a uniform concentration (Figures 5c and 8d, h). There were no statistically significant differences in distance from the peak between the naïve animals and the worms adapted to the low nicotine concentration (Figure 5c, F-NA0 vs. F-NA0.01).

Due to the nature of the behavior of the nematode *C. elegans*, the distribution of all the measured values was not normal. Therefore, for this type of distribution, an appropriate measure of central tendency is the median. Nonparametric tests more easily demonstrate the existence of a significant statistical difference, especially when a large amount of data is available. In our study, a single experiment provides 7200 data, because records were taken every 0.5 s during the 3600 s of the experiment. Additionally, the experiments were repeated 5 times, to increase the pool of results. Thus most of the experimental variants differed significantly from one other. In our view, more biologically important are the results which do not differ in a statistically significant way, despite the distinctly different experimental conditions.

*Caenorhabditis elegans* chemotaxes to bacteria, its natural food source, by following both water-soluble and volatile cues. Because of this difference in sensitivity, and because the diffusion of small molecules through air is much more rapid than through water, it is likely that volatile odors are used first for long-range chemotaxis, and later water-soluble attractants are used for short-range chemotaxis (Bargmann, 2006). Many studies have shown that the absence or presence of food markedly influences the average speed of wild-type worms (de Bono and Bargmann, 1998; Ramot et al., 2008). Sawin et al. (2000) reported that the feeding status (well-fed or starved) as well as the presence or absence of food (bacteria) affects the rate of locomotion of *C. elegans*. Sawin et al. (2000) reported a decrease in the locomotory rates of well-fed adult *C. elegans* on a bacterial lawn (food) compared with those on plates lacking bacteria, and defined this response as the “basal slowing response“. Nicotine exposure alters behaviors in *C. elegans*, including pharyngeal pumping, which disturbs nutrition (Matta et al., 2007). Pharynges that have been dissected from wild-type worms hypercontract in 0.1 mM nicotine (McKay et al., 2004). Nematodes in the presence of high concentrations of nicotine have smaller body size (Figure 4) probably because they eat less, as compared to control nematodes, thereby releasing less serotonin, and do not show enhanced slowing on food (Figures 5a and 6k, l). The neurosecretory-motor neurons that have sensory endings in the lumen of the pharynx (which thus might sense food) also synthesized serotonin (Chase and Koelle, 2007). Serotonin is required for the so-called “enhanced slowing response” (Sawin et al., 2000). *C. elegans* individuals adapted to the high nicotine concentration were smaller than those in the control, probably because in the presence of food their serotonergic neurons did not release an increased amount of serotonin, which did not inhibit the motor circuit to a greater extent than in the basal slowing response to effect the enhanced slowing response (Figure 5a F+nA0 vs. F+nA1). Without such serotonin release, the enhanced slowing response could not occur. Thus on the plate with food, the worms adapted to the high nicotine concentration moved faster than the control worms. The animals slow their locomotion rate dramatically when they encounter a bacterial lawn (Figures 5a and 6c, d, g, h) and, as mentioned above, the enhanced slowing response requires serotonin (Sawin et al., 2000). Serotonin signaling thus provides a mechanism to ensure that animals absolutely do not leave a food source once they have encountered it (Chase and Koelle, 2007). That is probably why we observed a very short distance from the start position for the control worms and a little larger distance for those adapted to the low concentration to nicotine (Figure 7c, g).

Sellings et al. (2013) were the first to reveal that nicotine serves as a primary motivating stimulus in *C. elegans*. Their research suggests that dopamine (DA) transmission is important in mediating the nicotine-motivated behaviors (Sellings et al., 2013). The presence of bacteria is perceived by mechanosensory stimulation via 8 dopaminergic sensory neurons (Sawin et al., 2000). The level of dopamine is higher in the presence of nicotine. We hypothesized that in nematodes adapted to high nicotine concentration (1 mM), the alkaloid mimics bacterial mechanosensory stimulation via sensors of dopaminergic neurons, thus nematodes eat less and have smaller body size (Figure 4a, F-nA0 vs. F-nA1). The dopamine signaling in well-fed animals encourages the animal to stay in the proximity of food but still may permit limited exploration for new or better food sources (Chase and Koelle, 2007). This is a likely explanation why the presence of nicotine encouraged nematodes to move away from the application site on the Petri dish (Figure 7d, h).

The effects of nicotine on locomotion vary according to dose and over time (Sobkowiak et al., 2011). In other animal models, acute administration of nicotine evokes also dual changes in locomotor activity (Matta et al., 2007). Innate chemosensory preferences are often encoded by sensory neurons that are specialized for attractive or avoidance behaviors. Tsunozaki et al. (2008) show that one olfactory neuron in *Caenorhabditis elegans*, AWC^ON^, has the potential to direct both attraction and repulsion. Attraction, the typical AWC^ON^ behavior, requires a receptor-like guanylate cyclase GCY-28 that acts in adults and localizes to AWC^ON^ axons (Tsunozaki et al., 2008). Interestingly, in our previous research we found GCY-28 only in naïve worms, which were distinctively attracted by nicotine. GCY-28 was absent in the protein complexes involved in response to low and high concentration of nicotine (Sobkowiak et al., 2016). This may suggest that in the presence of nicotine we observed the same kind of switching by presence or absence of food in control nematodes and worms adapted to low nicotine concentration (Figure 5d).

Nicotine induces profound behavioral responses in *C. elegans* that mimic those observed in mammals. The genes and pathways regulating nicotine dependence in mammals are functionally conserved in *C. elegans*, including nicotinic acetylcholine receptors (nAChRs, the molecular target of nicotine) and serotonin and dopamine-mediated neurotransmission. In this study, the exposure started from hatching and lasted 55 h, i.e. at least 3/4 of the *C. elegans* life cycle. Sellings et al. (2013) demonstrated, and we confirmed this finding, that *C. elegans* exhibits motivational behavior patterns towards nicotine, similar to those observed in mammalian models. The motivational behavior pattern is modulated, as we show, by food.

We revealed that nicotine can reduce worm body size (Figure 4). The anorectic effects of smoking have been well documented in human subjects, and the principal reason cited by female teenagers for why they smoke is weight control. On average, smokers weigh 5 kg less than nonsmokers and have significantly lower body mass index than nonsmokers. Similarly, nicotine decreases feeding in animal models, suggesting that nicotine in tobacco is important for the effects of smoking on appetite (Picciotto and Kenny, 2013).

Our results confirm that *C. elegans* may serve as a useful model organism for nicotine-motivated and food-motivated behaviors that could help in understanding the mechanisms underlying the anorectic effects of taking nicotine. Thus research on *C. elegans* may facilitate the development of novel treatments to help with smoking cessation and with preventing or treating obesity.

## Acknowledgments

We would like to thank the Caenorhabditis Genetics Center, which is funded by the NIH National Center for Research Resources (NCRR), for supplying the nematode strain used in this research. We thank also Sylwia Ufnalska, an author's editor, for help in improving the manuscript. This study was supported by funds from the Department of Cell Biology, Faculty of Biology, Adam Mickiewicz University (Poznań, Poland).

## Notes

**Competing interests:** The authors have declared that no competing interests exist.

